# Meta-Align: A Novel HMM-based Algorithm for Pairwise Alignment of Error-Prone Sequencing Reads

**DOI:** 10.1101/2020.05.11.087676

**Authors:** Kentaro Tomii, Shravan Kumar, Degui Zhi, Steven E. Brenner

## Abstract

**Background:** Insertion and deletion sequencing errors are relatively common in next-generation sequencing data and produce long stretches of mistranslated sequence. These frameshifting errors can cause very serious damages to downstream data analysis of reads. However, it is possible to obtain more precise alignment of DNA sequences by taking into account both coding frame and sequencing errors estimated by quality scores.

**Results:** Here we designed and proposed a novel hidden Markov model (HMM)-based pairwise alignment algorithm, Meta-Align, that aligns DNA sequences in the protein space, incorporating quality scores from the DNA sequences and allowing frameshifts caused by insertions and deletions. Our model is based on both an HMM transducer of a pair HMM and profile HMMs for all possible amino acid pairs. A Viterbi algorithm over our model produces the optimal alignment of a pair of metagenomic reads taking into account all possible translating frames and gap penalties in both the protein space and the DNA space. To reduce the sheer number of states of this model, we also derived and implemented a computationally feasible model, leveraging the degeneracy of the genetic code. In a benchmark test on a diverse set of simulated reads based on BAliBASE we show that Meta-Align outperforms TBLASTX which compares the six-frame translations of a nucleotide query sequence against the six-frame translations of a nucleotide sequence database using the BLAST algorithm. We also demonstrate the effects of incorporating quality scores on Meta-Align.

**Conclusions:** Meta-Align will be particularly effective when applied to error-prone DNA sequences. The package of our software can be downloaded at https://github.com/shravan-repos/Metaalign.

## Background

Metagenomics sequencing projects, the application of sequencing to environmental samples, have accumulated large numbers of DNA sequences and revealed millions of new protein coding genes [1]. However, the read coverage of typical metagenomic data results in many incomplete protein sequences and propagates sequencing error rates to gene predictions. With respect to error rates, next-generation sequencing (NGS) technologies, such as pyrosequencing, ion semiconductor sequencing and single molecule real time (SMRT) sequencing, used in metagenomics have relatively higher rates of insertion and deletion (indel) sequencing errors (i.e., too few or too many bases called) [2]. When present in a protein coding region an indel error causes a frameshift in the resulting gene, so that subsequent amino acids and the termination codon are incorrectly predicted. A situation in which the frameshifted sequence does not terminate prematurely is even worse, since the resulting scrambled amino acid sequence serves only to introduce noise. In combination, these effects mean that frameshift errors can produce very serious difficulties in any downstream analysis of reads. Frameshifts will be common in metagenomic sequence data because a large fraction of organism’s genomes will be covered only with single- or a few-pass reads. In such circumstances, improving computational tools is required for the efficient analysis of those metagenomic data.

When nucleotide sequences are available, a more precise alignment is possible that takes into account both coding frame and frameshifts. It is not desirable to simply align at the DNA level, since it is well known that for distant homology detection, it is far more effective to compare protein sequences than their coding DNA sequences (see e.g., [3]) because selection takes place at the level of the proteins. Thus, physicochemical similarities between amino acid residues are far more important in determining homology at moderate distances than the match of the aligned nucleotides that encode them. Furthermore, standard nucleotide gap penalties increase with length, with the result that a gap of length one will be assigned a smaller penalty than a gap of length three. However, in a coding region the reverse should be true, as a frameshift (implied by a gap of length one) is much less probable than a single amino acid deletion (gap of three nucleotides). In a typical nucleotide-peptide alignment framework such as BLASTX [4] which searches protein database using a translated nucleotide query, nucleotide sequences are translated to yield amino acids. However, this technique has several drawbacks when working with error-prone DNA sequence. In particular, it assumes that the reading frame is known perfectly throughout the whole sequence. In the presence of indels, this is not the case. For this reason, several groups have proposed variants of the Smith-Waterman-Gotoh algorithm [5, 6] to not only align DNA sequences scoring a codon at a time with an amino acid substitution matrix, but also allow frameshifts in the sequence [7-9]. For instance, Irie et al. devised the nucleotide sequence alignment method which consider amino acid similarity and indels at the both DNA and protein levels, although the method only allows an insertion or a deletion for a codon at the DNA level [9].

Another major issue in the analysis of NGS reads is incorporating quality scores from reads. By exploiting per-base quality scores, such as the Phred Q-score [10, 11], we anticipate further improvement in frameshift-permitting sequence alignment. For example, an alignment of the codon TGG with TAG has one nucleotide difference, but implies the low-probability substitution of Tryptophan with a stop codon. However, if the A in TAG has a poor Q-score, this increases the probability that the A is a mis-called G and that the Tryptophan is conserved.

Incorporating quality information of bases to weight mismatches in the DNA sequence alignments is done for the comparison of EST sequences [12], and for mapping reads to a genomic sequence with use of multi- (3 or 4) dimensional substitution matrix [13].

However, sequence similarities at the protein level are not taken into account in both cases. To the best of our knowledge, there is no such method that would simultaneously consider both amino acid substitution probabilities implied by aligned pairs of codons and the quality scores for called DNA bases to align coding regions of DNA sequences. We therefore designed and implemented a novel pairwise alignment algorithm, called Meta-Align, that allows for aligning error-prone reads in amino acid (protein) space, incorporating quality scores from the DNA sequences, and tolerating frameshifts caused by insertions and deletions. The alignment algorithm is capable of accounting for the frameshifts, substitutions, and indels, and obtaining the most likely alignment of the amino acids produced by matching pairs of triplets, that are predicted codons in each source DNA sequence. Each triplet may contain any numbers of gaps in its constituent DNA sequence, thereby seamlessly accommodating frameshift errors into the alignment process. In particular, unlike frameshift matrices, this type of algorithm has the ability to extend alignments across the point of a frameshift, which we expect to increase sensitivity when appropriately modified to compute local alignments. We anticipate that our algorithm will be particularly effective when applied to error-prone DNA sequences.

## Methods

### Overview of the model

Our model is partly inspired by the comparative genomics gene finding HMM GeneWise [14], which uses an HMM transducer framework that combines a gene prediction HMM and a pair HMM homology model. In general, a pair HMM of two output sequences can be viewed as a transducer that translates one output sequence to the other output sequence. One can concatenate several transducers into a pipeline that translates one output sequence to the other through a series of intermediate translations. The transducer model has been used to various computational biology problems including pairwise gene finding [14] and phylogenetics contexts [15]. In our model, we consider the similarity between two input metagenomics read sequences through 4 intermediate sequences via 5 pairwise transducers (Figure 1). The full model can be specified by spelling out the transducer by applying the automata theory [16].

**Figure 1.**
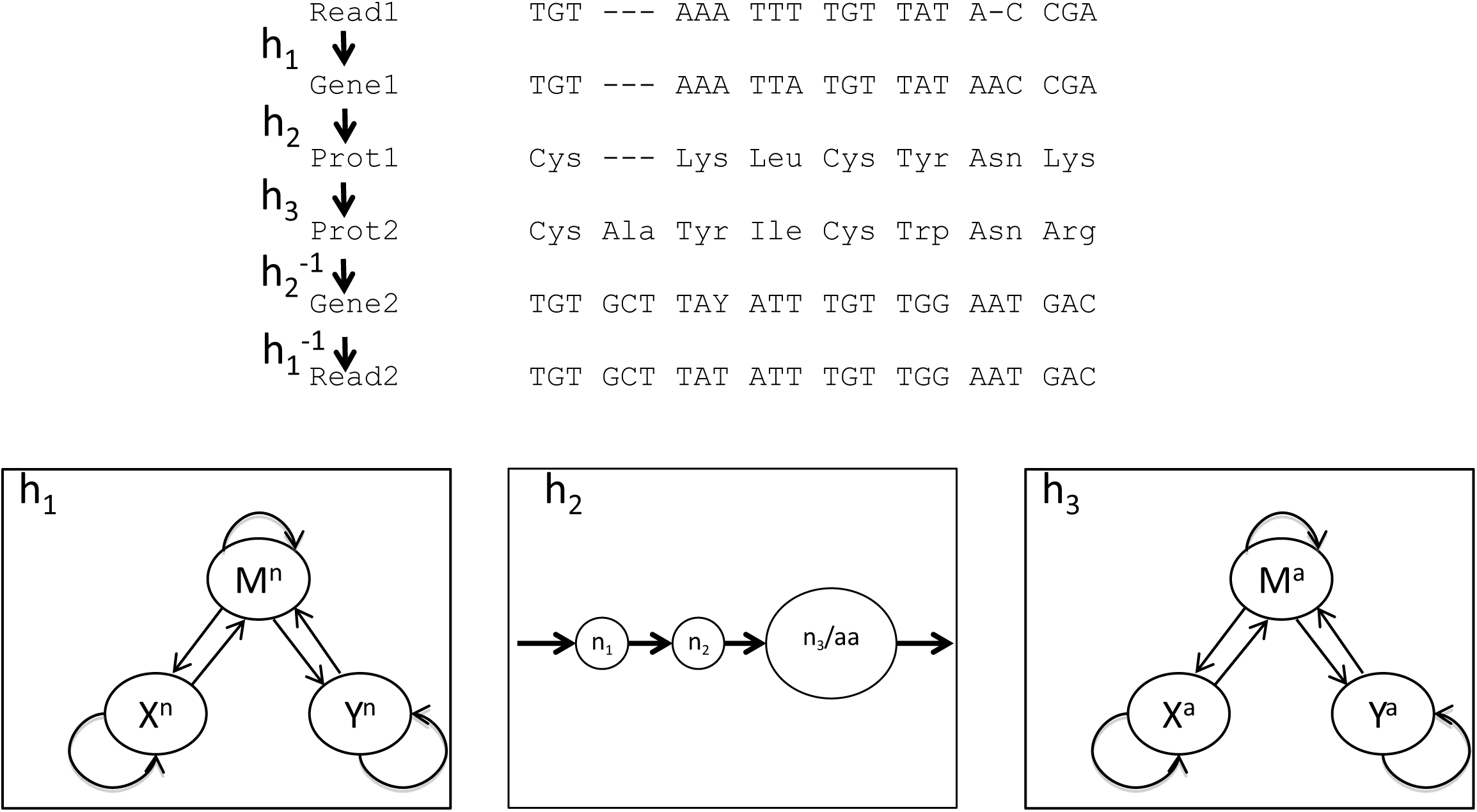
Transducer HMM of Meta-Align. Meta-Align consists of five pair HMMs, h_1_, h_2_, h_3_, h_2_^-1^, and h_1_^-1^·h_1_, and h_1_^-1^ model the sequencing process, where state M^n^ is the sequencing read out, with emission probability encoding sequencing substitution errors; states X^n^ and Y^n^ corresponds to sequencing insertion and deletion errors. h_2_ and h_2_^-1^ are 2^nd^ order HMMs corresponding to the genetic code, where states n_1_ and n_2_ emit the nucleotides at the first two codon positions, and the state n_3_ /aa emits the nucleotide at the last codon position and the encoded amino acid, depending on the previous two codon positions. The central HMM h_3_ models the protein sequence alignment, where state M^a^ is the match state parameterized by an amino acid substitution matrix, and states X^a^ and Y^a^ corresponds to insertion and deletion.

We have developed our model based on both an HMM transducer of a pair HMM and profile HMMs so that the model can align two reads by considering simultaneously both similarity at the protein level and quality information of bases at the nucleotide level. The schematic representation of our HMM topology is shown in Figure 2. Our model consists of three parts, i.e. the Match/Mismatch, Insertion and Deletion moieties. These moieties correspond to the Match/Mismatch state, Insertion state, and Deletion state, respectively, in a standard three-state pair HMM for pairwise sequence alignment. Profile HMM units, designed to predict a codon pair in the true sequences within each moiety, are intended to incorporate quality information of reads to obtain most likely alignments.

**Figure 2.**
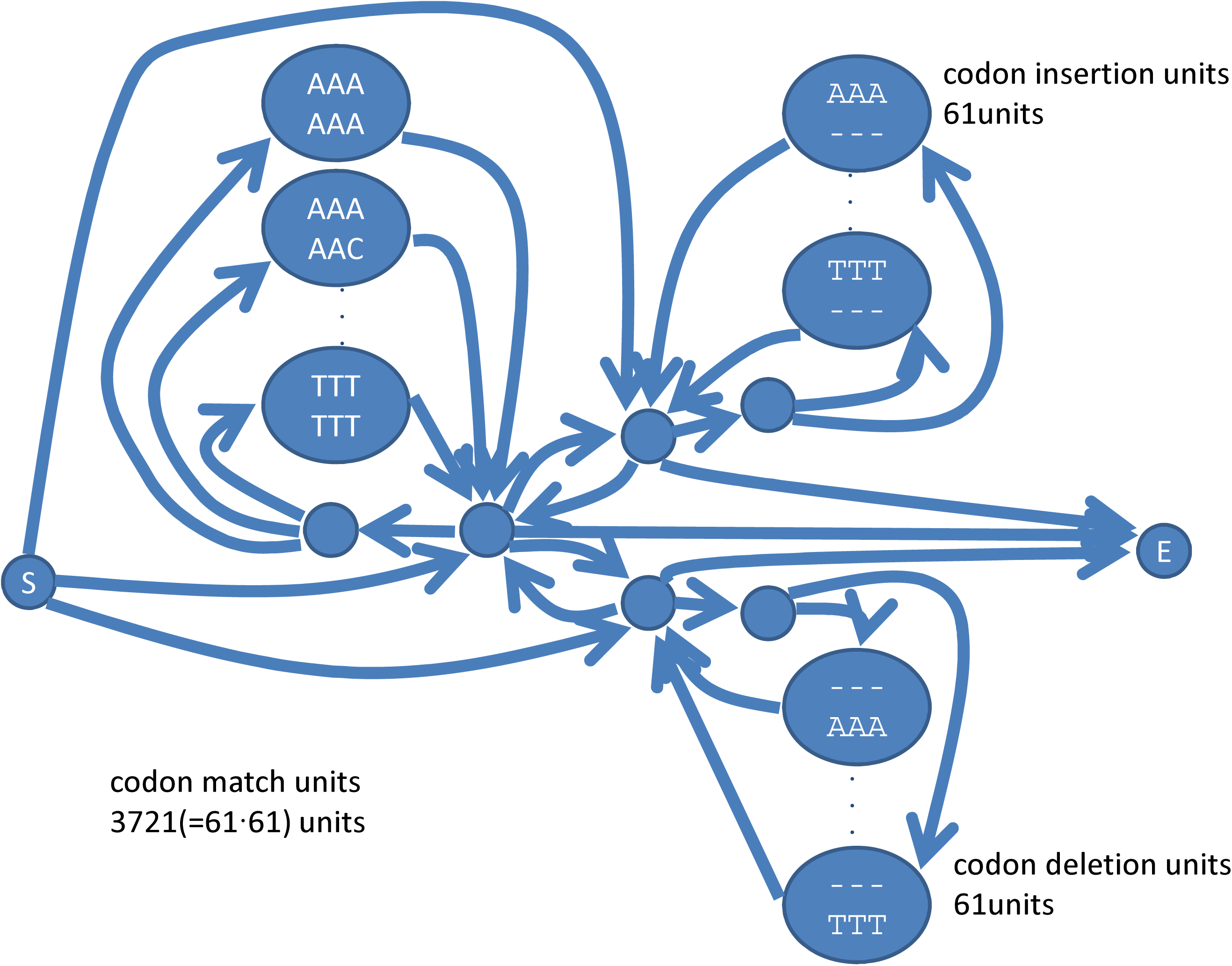
Schematic of the *full* HMM of Meta-Align. See text for a detailed description.

### Algorithms

#### Elaborate profile HMMs for DNA alignment and frameshift detection

Let the two nucleotide sequences (i.e., reads) be *X* = *x*_*1*_ *x*_*2*_ …*x*_*n*_ and *Y* = *y*_*1*_ *y*_*2*_ …*y*_*m*_, and their quality scores be *Q* = *q*_*1*_, *q*_*2*_, …, *q*_*n*_ and *R* = *r*_*1*_, *r*_*2*_, …, *r*_*m*_, respectively. Three (i.e. Match/Mismatch, Insertion and Deletion) moieties in our model is used to align nucleotide sequences, find sequencing errors, and calculate the probability of alignments based on the model of a codon in the *true* DNA sequence.

The Match/Mismatch moiety consists of 3721(=61×61) codon Match/Mismatch unit (CMU) for all possible amino acid pairs, and the Insertion and Deletion moieties have 61 units, respectively. We named this model as the *full* model. In a CMU each triplet (a predicted codon in the *true* sequence) may contain any numbers of gaps in its constituent DNA sequence, thereby seamlessly accommodating frameshift errors into the alignment process. As an example, we show a CMU, based on an elaborate profile HMM (Figure 3), which contains 22 states and 108 transitions. A CMU has six types of states, i.e., match *M*, insertion *I*_*x*_, deletion *D*_*x*_, insertion *I*_*y*_, deletion *D*_*y*_, and *silent* double deletion *D*_*d*_ states, in addition to the *Begin* and *End* states. The recursion equations are as follows:

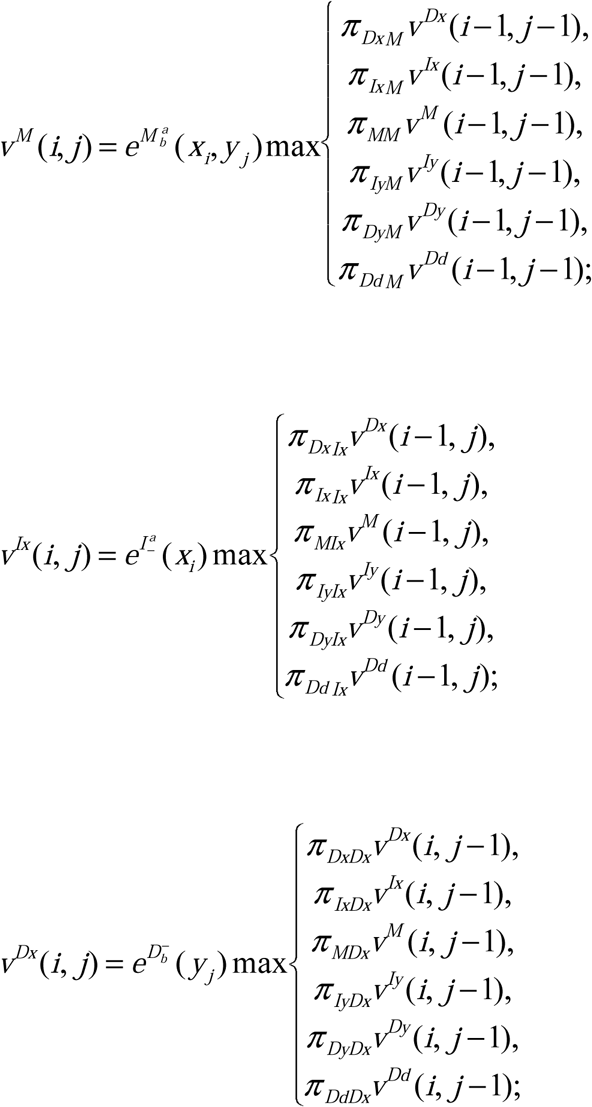

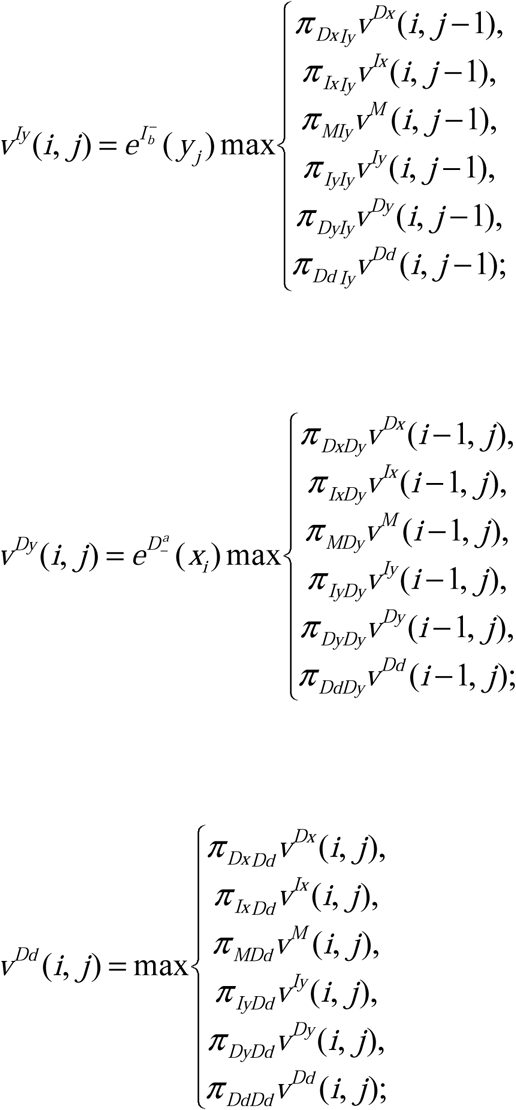

where *e* indicates an emission probability depending on quality scores. For instance, the emission probability of a match state,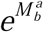 is calculated as:

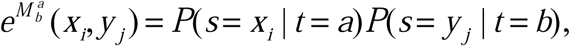

where *a* and *b* indicate given types of bases, i.e., *a, b* = {A, C, G, T}. *P(s=x*_*i*_ |*t=a)* is the conditional probability of observing a called base *x*_*i*_ in a read, given a *true* base *a* in a genome (*P(s=y*_*j*_ |*t=b)*, in turn). These probabilities are given by

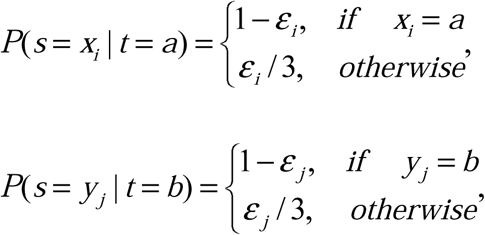

where *ε*_*i*_ and *ε*_*j*_, which represent the probabilities of base call errors, are calculated based on quality scores according to the well known Phred Q-score definition: *ε*_*i*_ = 10^−*qi*/10^ and *ε*_*j*_ = 10^*-rj*/10^. Thus, the larger the quality score, the larger the probability of the case when *x*_*i*_ =*a*, which means the *correct* calling probability, given a *true* base *a* (*y*_*j*_ = *b*, in turn). We give equal probabilities for the three rest cases (i.e., *x* _*i*_ ≠ *a*), because we assume unbiased base call error for simplicity. Similarly, the emission probability 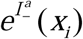 of an insertion state *I*_x_ is defined as 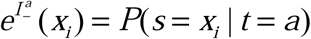 *π*s indicate transition probabilities between two states in a CMU. We treat indels under the stochastically simplifying assumption that these are computed from the error frequency. For instance, we define the transition probability *π*_*_ *Ix* to an insertion state *I*_*x*_ as *π*_* *Ix*_ =*ε*_*i*_ *P(Ins)*, where ‘*’ indicates any states in a CMU, i.e., * = *I*_*x*_, *D*_*x*_, *M, I*_*y*_, *D*_*y*_, *D*_*d*_ (and the *begin* state). Thus, the transition probability to a match state is: *π*_* *M*_ =1-{*ε*_*i*_ *P(Ins)*+*ε*_*i*_ *P(Del)*+*ε*_*j*_ *P(Ins)*+*ε*_*j*_ *P(Del)*+*ε*_*i*_ *P(Del)ε*_*j*_ *P(Del)*}. Precisely speaking, adding the last term, *ε*_*i*_ *P(Del)ε*_*j*_ *P(Del)* leads to the overestimation of error probability. However, because the term has usually extremely small probability, we adopted this approximation for simplicity. The values, such as given in Table 1 of [2], can be adapted to *P(Ins)*, which represents the rate of insertion sequencing errors (*P(Del)*, in turn). The probabilities to the *End* state are 1, i.e., *π*_* *End*_ =1, where *= *I*_*x*_, *D*_*x*_, *M, I*_*y*_, *D*_*y*_, and *D*_*d*_.

**Table 1.** Comparison of the performances of Meta-Align and TBLASTX using simulated data sets. Each line reports the summary of 100 randomly sampled read pairs using a particular parameter setting (mutation rate, indel count, and codon bias). The *FD* and *FM* scores are calculated based on the structurally aligned amino acid sequences from the BAliBASE. P-values of one-sided paired t-tests were used to evaluate the statistical significance of the difference between Meta-Align and TBLASTX.

| Mutation<br>rate, $\alpha$ | InDel<br>count | Condon<br>bias, $\beta$ | FD | FD | FM | FM | P-Value<br>(FD) | P-Value<br>(FM) |
| --- | --- | --- | --- | --- | --- | --- | --- | --- |
|  |  |  | Meta-Align | TBLASTX | Meta-Align | TBLASTX |  |  |
| 0.005 | 0 | 0 | 0.715 | 0.384 | 0.727 | 0.549 | 2.89E-22 | 6.75E-07 |
| 0.005 | 1 | 0 | 0.683 | 0.305 | 0.695 | 0.556 | 1.48E-30 | 2.44E-05 |
| 0.005 | 2 | 0 | 0.623 | 0.268 | 0.638 | 0.516 | 4.53E-27 | 2.80E-04 |
| 0.02 | 0 | 0 | 0.675 | 0.362 | 0.689 | 0.523 | 1.95E-22 | 4.64E-07 |
| 0.02 | 1 | 0 | 0.649 | 0.282 | 0.662 | 0.487 | 6.80E-26 | 2.21E-07 |
| 0.02 | 2 | 0 | 0.597 | 0.249 | 0.609 | 0.497 | 2.12E-23 | 1.33E-03 |
| 0.05 | 0 | 0 | 0.635 | 0.313 | 0.654 | 0.465 | 1.87E-21 | 6.28E-07 |
| 0.05 | 1 | 0 | 0.576 | 0.222 | 0.593 | 0.448 | 4.88E-23 | 1.33E-04 |
| 0.05 | 2 | 0 | 0.564 | 0.210 | 0.582 | 0.441 | 4.74E-24 | 2.00E-04 |
| 0.005 | 0 | 0.5 | 0.701 | 0.397 | 0.713 | 0.561 | 1.68E-22 | 2.07E-06 |
| 0.005 | 1 | 0.5 | 0.685 | 0.313 | 0.697 | 0.568 | 1.45E-30 | 2.94E-05 |
| 0.005 | 2 | 0.5 | 0.651 | 0.265 | 0.662 | 0.530 | 1.15E-32 | 1.28E-04 |
| 0.02 | 0 | 0.5 | 0.684 | 0.359 | 0.699 | 0.477 | 1.06E-21 | 6.40E-10 |
| 0.02 | 1 | 0.5 | 0.651 | 0.260 | 0.667 | 0.472 | 2.67E-28 | 2.17E-07 |
| 0.02 | 2 | 0.5 | 0.630 | 0.257 | 0.640 | 0.501 | 8.41E-28 | 3.38E-05 |
| 0.05 | 0 | 0.5 | 0.652 | 0.294 | 0.670 | 0.445 | 1.83E-23 | 1.11E-08 |
| 0.05 | 1 | 0.5 | 0.614 | 0.257 | 0.631 | 0.456 | 1.25E-26 | 1.09E-06 |
| 0.05 | 2 | 0.5 | 0.584 | 0.229 | 0.598 | 0.509 | 8.34E-27 | 5.57E-03 |
| 0.005 | 0 | 1 | 0.572 | 0.384 | 0.584 | 0.559 | 2.99E-12 | 2.39E-01 |
| 0.005 | 1 | 1 | 0.557 | 0.281 | 0.566 | 0.511 | 3.18E-21 | 3.53E-02 |
| 0.005 | 2 | 1 | 0.547 | 0.232 | 0.557 | 0.497 | 5.40E-26 | 4.17E-02 |
| 0.02 | 0 | 1 | 0.549 | 0.370 | 0.562 | 0.530 | 4.46E-14 | 1.56E-01 |
| 0.02 | 1 | 1 | 0.551 | 0.285 | 0.561 | 0.552 | 2.00E-18 | 3.88E-01 |
| 0.02 | 2 | 1 | 0.513 | 0.241 | 0.524 | 0.534 | 1.30E-19 | 3.86E-01 |
| 0.05 | 0 | 1 | 0.536 | 0.336 | 0.551 | 0.527 | 1.68E-10 | 2.67E-01 |
| 0.05 | 1 | 1 | 0.509 | 0.240 | 0.521 | 0.498 | 3.88E-20 | 2.52E-01 |
| 0.05 | 2 | 1 | 0.492 | 0.185 | 0.504 | 0.419 | 5.92E-22 | 1.07E-02 |

**Figure 3.**
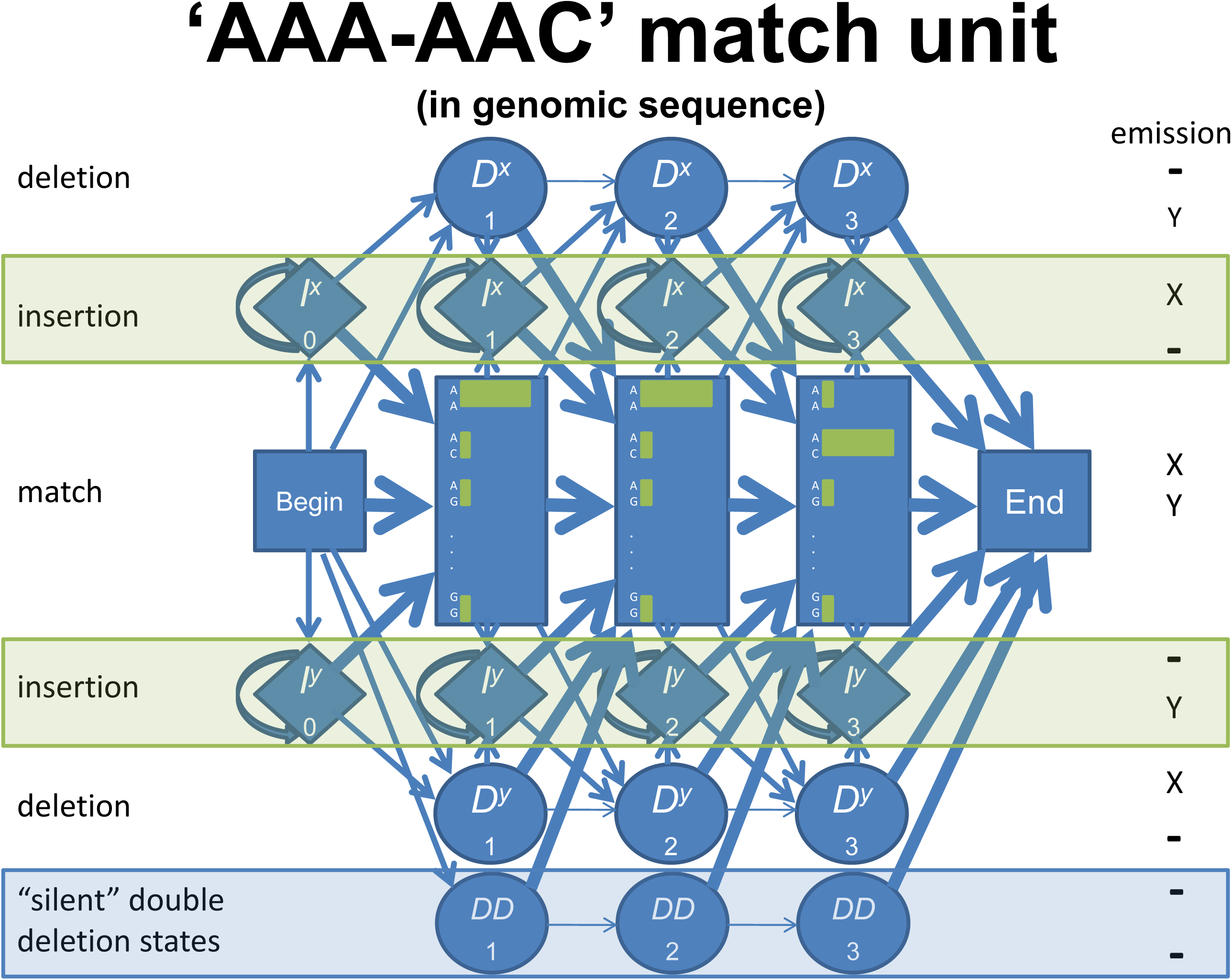
A codon Match/Mismatch unit (CMU) in the *full* HMM. A CMU has six types of states, i.e., match states that emit aligned pairs of *x*_*i*_ and *y*_*j*_, insertion (happened in *X*) states that emit *x*_*i*_ and a gap, deletion (happened in *X*) states that emit a gap and *y*_*j*_, insertion (happened in *Y*) states that emit a gap and *y*_*j*_, deletion (happened in *Y*) states that emit *x*_*i*_ and a gap, and *silent* double deletion states that emit nothing.

We note that the occurrence of indel errors is non-random in sequences, for example occurring in homopolymers by technologies such as 454 pyrosequencing [17] and ion semiconductor sequencing [18]. Here we do not account for this directly, as it creates a problem of scalability to our model. Rather, we will rely on base callers that have intimate knowledge of the sequencing technology to provide accurate quality scores. We anticipate that a wide range of parameters for frameshift transitions based on quality scores will yield satisfactory results, which we evaluated as below.

#### Pair HMM for alignment at the protein level

An extended pair HMM in our model is used for evaluating the similarity of two reads at the protein level. The model is based on a standard three-state pair HMM for pairwise alignment of amino acid sequences, and is comprised of six states, *M’, X’*, and *Y’*, and their respective dummy states, in addition to three moieties described above, and the *begin* and *end* states. *X’* and *Y’* correspond to the gap (Insertion and Deletion) states and *M’* corresponds to the Match/Mismatch state in protein space. We can obtain the optimal alignment of two reads by using the Viterbi algorithm for pair HMM. The recursion equations are:

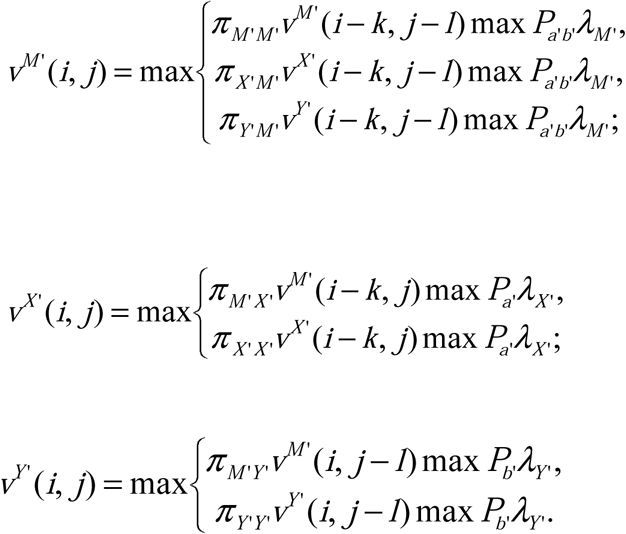

Here *λ*_*M’*_, *λ*_*X’*_, and *λ*_*Y’*_ indicate respectively the probability calculated by a CMU, that consists of an elaborate profile HMM, in the Match/Mismatch, Insertion, and Deletion moiety described above. *P*_*a’b’*_ denotes the target frequency, derived from any appropriate substitution matrix [19], for the amino acid pair, *a’* and *b’. P*_*a’*_ and *P*_*b’*_ are the amino acid compositions of *a’* and *b’*, respectively. *k* and *l* are arbitrary positive integers. *k* = *l* = 3, if there is no gap in a triplet. Again, any numbers of gaps are considered in a codon by a CMU. *π*s correspond to transition probabilities between any pair of states among three states in a standard three-state pair HMM for pairwise alignment of amino acid sequences. The transition probability from *M’* to its dummy state is: *π*_*M’M’*_ =1-2*δ*-*τ*, and the ones from *M’* to *X’* or *Y’* are: *π*_*M’X’*_ =*π*_*M’Y’*_ =*δ*. The transition probabilities from *X’* or *Y’* to *M’* are: *π*_*X’M’*_ =*π*_*Y’M’*_ =1-*ε’*-*τ*. The transition probabilities from *X’* or *Y’* to their respective dummy states are: *π*_*X’X’*_ =*π*_*Y’Y’*_ =*ε’*. These parameters, *δ* and *ε’*, associated with gap opening and extension, and *τ* are defined as usual manner in a standard pair HMM for pairwise alignment. In our implementation, we choose the following values for the parameters: *δ* = 0.20, *τ* = 0.02, *ε’* = 0.49. Herewith, the alignment algorithm generates the optimal alignment of the amino acids produced by matching pairs of triplets from each source DNA sequence.

To obtain the transition probabilities from dummy states to CMUs in each moiety, we combine target frequencies with codon usage frequencies in a genome. For instance, the transition probability from the dummy state of *M’* to the AAA-AAG CMU is defined as the product of the target frequency for the Lysine-Lysine pair and the joint probability of the codon usage frequencies of AAA-AAG pair. The frequencies appeared in [20] can be used. The alignment algorithm computes the score for matching two amino acid distributions by using the target frequencies, weighted by the joint probability of each codon pair. Therefore, our model is completely consistent with a standard three-state pair HMM for pairwise sequence alignment, when there are no uncertainties in given reads.

#### Degenerative model

Because of the sheer number of states of the *full* model described above, it is not computationally feasible. We thus derive a degenerative model which is essentially equivalent to the full model, while reducing the number of states dramatically. The reduction of model is based on the degeneracy of the genetic code: the amino acid is mostly determined by the first two codon position, while the third codon position allowed to “wobble”. Our idea is to collapse the 16 CMUs in the *full* model representing matching between four AAX codons, including AAA, AAC, AAG, and AAT, and four TTX codons, including TTA, TTC, TTG, and TTT into one degenerative CMU (Figure 4). We recognize that all 16 CMUs share the HMM model structure relevant to the first two codon positions, so we keep this in the degenerative CMU. In the third codon position we use a single AAX-TTX third codon match state, AAX-TTX-M3, with an elaborated 4-by-4 emission probability table.

**Figure 4.**
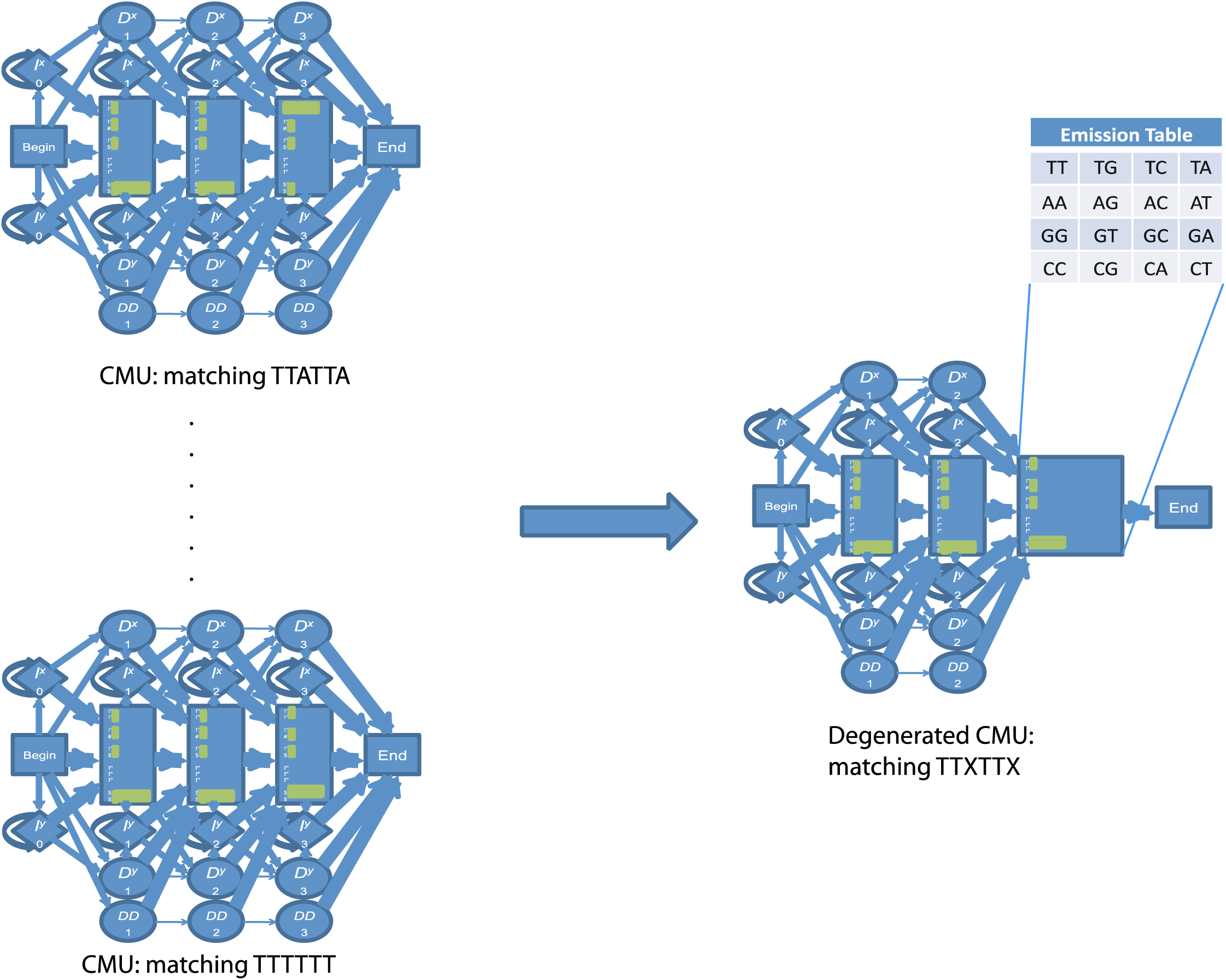
Schematic of collapsing the 16 CMUs in the *full* model into a single CMU in the degenerated model. See text for a detailed description.

The emission probabilities are adjusted so it approximates the emission-transition probabilities of the relevant path in the *full* model. For example, since AAX translates into Lysine (AAA and AAG) and Asparagine (AAC and AAT), and TTX translates into Leucine (TTA and TTG) and Phenylalanine (TTC and TTT), we set the emission probability table of AAX-TTX-M3 as the following:

*P*_*d*_*(emit A/A)=P*_*f*_*(codon-match AAA-TTA)*,

…

*P*_*d*_*(emit T/T)=P*_*f*_*(codon-match AAT-TTT)*,

where *P*_*d*_ is a probability in the degenerated model and *P*_*f*_ is a probability in the *full* model. Using the redundancy in the third position we reduce 3721 CMUs in the *full* model to 256 (=16×16) degenerated CMUs. Moreover, the CMUs in the *full* model have 22 states, while the CMUs in the degenerated model have 17 states, due to the simplification at the third codon position. Overall, the number of states in the generated model reduced 14.27-fold (92%) compared to the *full* model.

We note that the degenerate model trades of completeness for computational feasibility. The *full* mode is capable of handling insertions in all three codon positions, while the degenerate model is capable of handling only insertions in the first two positions. However, this loss of capability is unlikely an issue in practice because that an alignment with an insertion at the third codon position can be approximated by a similarly scored alignment with an insertion at the nearby first or second codon position.

#### Implementation

We implemented the algorithm in C++ as an extension of the existing HMMOC package [21]. The output alignment is calculated as the most probable path through the HMM determined by the Viterbi algorithm. A software package can be downloaded at https://github.com/shravan-repos/Metaalign.

### Simulation protocol for evaluation

BAliBASE is a collection of expert-annotated structurally alignment of protein sequences [22], and is commonly used for benchmarking sequence alignment algorithms. In the present work, subsets of BAliBASE with varying sequence identities and lengths were considered. We chose the virtual read length to be about 200 nt, the typical length of a 454 pyrosequencing read, in our simulation. This corresponds to 70 amino acids after reverse translation.

The first step of our simulation is sampling of pair of aligned read sequences as following: A multiple sequence alignment is randomly selected from the BAliBASE and a pair of sequences of the multiple alignment is randomly chosen. If the selected sequences have more than 70 amino acids, random trimming is done and an interval consisting of exactly 70 amino acids is selected. However, when short sequences containing less than 70 amino acids are chosen, we used those sequences without any additional trimming or padding. The second step is an *in silico* reverse-transcription from amino acid sequences to *genomic* DNA sequences. The genetic code has degeneracy. One amino acid can be encoded by multiple codons. In a species one codon may be favored than others, creating codon bias. In our simulation, we define a parameter for codon bias, *β*, as following. If an amino acid has *k* codons, the probability of the first codon (in alphabetic order) *C*_*1*_ is *P(C*_*1*_ *)=1-β(k-1)/k*; and the probability of any other codon is *P(C*_*j*_ *)=β/k*, for *j*>1. Therefore, when *β*=1, the reverse translation goes to the first codon deterministically; when *β*=0, all codons receive equal probabilities of *β/k*. In reality most species would have a codon bias in between. We consider *β*=0, 0.5, 1 in our simulation.

The third step of our simulation is to introduce mutations and sequencing errors during the generation of read sequences from the *genomic* DNA sequences. These mutations can be thought as the combined effect of additional evolutionary mutations (nonsynonymous and synonymous), and sequencing errors. The sequencing error rate of most sequencing platforms are around 1(∼2) %, with no preference of codon positions, while the evolutionary mutations typically are more concentrated on synonymous mutations or nonsynonymous mutations to similar amino acids than nonsynonymous mutations to dissimilar amino acids. In our simulation we do not intend to separate these two effects. Instead, we use a combined substitution rate, α, and vary it to be 0.5%, 2%, and 5%. In addition, we introduce single nucleotide insertions and deletions. These indels lead to frameshifts, which can drastically distort the translated amino acid sequences and thus create major challenge for aligning metagenomics reads. This is particularly a challenge for aligning reads from sequencing platforms prone to indels, such as 454 pyrosequencing. We test scenarios of having 0, 1, and 2 insertions.

It should be noted that since our pairwise alignment of read sequences is extracted from the BAliBASE multiple sequence alignments, the pairwise alignment derived from the BAliBASE may not be the alignment that maximizes a predefined score. We still use them as reference alignments on the bases that they represent *bona fide* structural alignments.

### Measuring metrics

Two types of scores are used to assess the quality of an alignment produced by Meta-Align: the Developer Score and the Modeler score, proposed by [23]. The Developer score *FD* is defined as the ratio of the number of correctly aligned residue pairs to the number of aligned residue pairs in the reference alignment. *FD* measures the sensitivity of the algorithm. The Modeler score *FM* is defined as the ratio of number of correctly aligned residues pairs to the total number of aligned residue pairs by the algorithm. *FM* measures the specificity of the algorithm.

The output of TBLASTX is essentially a list of local alignments. These alignments can represent overlapping regions, in different frames or strands. This contrasts with the output of Meta-Align, which is a global alignment. For the sake of comparison, we only consider the highest scoring region of TBLASTX. Therefore, it is likely that Meta-Align generates a longer aligned region (higher *FD*) than TBLASTX. What is more informative is whether Meta-Align also achieves higher specificities, i.e., offering higher *FM* scores than TBLASTX.

## Results

### Comparison of Meta-Align versus TBLASTX

We generated 100 pairs of virtual reads for each combination of codon bias, substitution rate, and indel count, totaling 2700 pairs of reads. Using the simulated data sets we compare the performance of Meta-Align and TBLASTX, which searches translated nucleotide database using a translated nucleotide query. As shown in Table 1, Meta-Align dominates TBLASTX, achieving significantly higher average *FD* scores across all parameter combinations and significantly higher average *FM* scores for almost all parameter combinations.

For “no codon bias” (*β*=0) and “some codon bias” (*β*=0.5) cases, Meta-Align outperforms TBLASTX for a wide margin in every one of the simulated sets. For “complete codon bias” (*β*=1) cases, Meta-Align has superior *FD* scores. However, the *FM* scores of Meta-Align and TBLASTX are similar, with Meta-Align marginally better. Moreover, there is no obvious effect of codon bias to the performance of TBLASTX.

Interestingly, while there is no obvious difference in performance of Meta-Align for “no codon bias” and “some codon bias” cases, Meta-Align has much worse performance for “complete codon bias” cases. This result indicates that, because the sequence identity is very low in our test cases, Meta-Align relies on the randomness signals in the codon choice to discern the coding frame and thus making the alignment. When complete codon bias is present, there is no signal for the coding frame and thus Meta-Align’s performance is degraded into one that is similar to TBLASTX.

For both Meta-Align and TBLASTX, two trends are observable. First, with increasing mutation rates, both *FD* and *FM* scores become lower. Second, with increasing indels, both *FD* and *FM* become lower. Interestingly, with increasing indels, the *FD* of TBLASTX decreases faster than the *FM* of TBLASTX. This is consistent with the idea that TBLASTX is a local alignment program and reports any local alignment above a significance threshold. With higher indel rates, TBLASTX finds shorter alignment (lower *FD* scores) but maintaining the alignment quality (*FM* scores). On the other hand, the HMM has advantage, because the HMM can compute the full probability that sequences *X* and *Y* could be generated by a given pair HMM; thus, a probabilistic measure can be introduced to help establish evolutionary relationships.

### Effect of amino acid level similarity on Meta-Align

To articulate the contribution of amino acid level similarity of our HMM model, we compare Meta-Align to a scaled down version of HMM model, HMM_NUC. The HMM_NUC model is the paired HMM model with only the nucleotide-level similarity and quality scores. Although HMM_NUC does a nucleotide level alignment, it still incorporates some amount of amino acid alignment information, not as complete as Meta-Align. Since it does not completely consider the sequence similarity at the amino acid level, it contains much smaller number of states and it runs drastically faster than Meta-Align.

We compare the performance of Meta-Align and HMM_NUC using the same simulated data set as in the above Meta-Align and TBLASTX comparison. As shown in Table 2, Meta-Align clearly outperforms HMM_NUC when mild codon bias present. Noticing this level of codon bias is the most relevant in real biological sequences, we believe that Meta-Align offer a higher quality alignment than HMM_NUC.

**Table 2.**
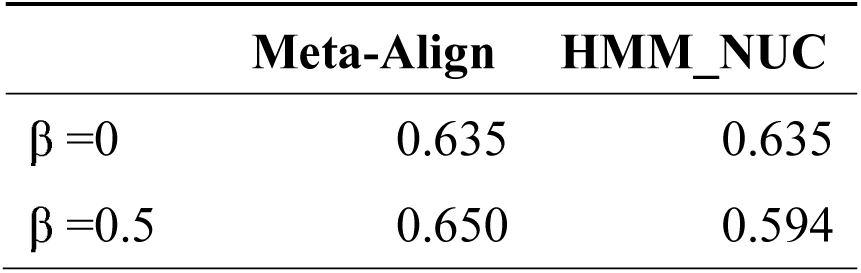
Comparison of Meta-Align versus HMM_NUC. The average *FD* scores for aligning 900 pairs of reads randomly sampled from BAliBASE alignments are presented.

|  | Meta-Align | HMM_NUC |
| --- | --- | --- |
| $\beta = 0$ | 0.635 | 0.635 |
| $\beta = 0.5$ | 0.650 | 0.594 |

### Effect of quality scores on Meta-Align

To study the effect of input quality scores on the performance of Meta-Align, we compare the result using real and fake quality scores. For a pair of *genomic* sequences reverse-transcribed from BAliBASE, we first generate quality scores for every nucleotide position, drawn from a uniform distribution from 5 to 30. Based on these quality scores we generate read sequences by introducing either a substitution (with a 70% probability) or an insertion (with a 30% probability). Subsequently, we run Meta-Align with two choices of quality score sequences: (i) the real quality score sequence based on which the sequences were generated, and (ii) a “fake” quality score sequence containing a constant value 20 at all nucleotide positions. Since Meta-Align takes into account the quality scores, it is expected to perform better when the real quality scores are used. We compared the results for 45 pairs of random sequences from BAliBASE and indeed Meta-Align with correct quality scores had a higher quality alignment at the amino acid level (Table 3).

**Table 3.** Comparison of Meta-Align using real quality scores versus using fake (flat) quality scores. The average results for 45 pairs of randomly chosen short sequences with varying sequence identity from BAliBASE are shown.

| Parameter | P-Value (Pair T-test) |
| --- | --- |
| FD (Amino Acid Sequence) | 0.02125 |
| FM (Amino Acid Sequence) | 0.02674 |
| FD (Nucleotide Sequence) | 0.3207 |
| FM (Nucleotide Sequence) | 0.5926 |

### The handling of different types of sequencing error

Traditionally there is only a single quality score indicating the probability of instrument reading error at a nucleotide position, and there is no way to distinguish between what type of error the instrument error might be. This could be a problem as it is known that some sequencing machines have specific error characteristics. For example, sequencers such as 454 and Ion Torrent are known to tend to make error in determining the length of a homopolymer run, resulting in higher indel error rates at such regions, while the probability of making a substitution reading error is similar to other regions.

We can specify different probabilities for different types of sequencing errors, accommodating characteristic sequencing errors resulting from different instruments. For this purpose, we can introduce two quality scores at each nucleotide position, *q*_*i*_, and *q*_*s*_, corresponding to the probabilities for indel errors and substitution errors, respectively. For a position with quality score *q*, we first assign the error probabilities according to the quality scores, *e*_*i*_ =*e*_*s*_ =10^−*q*/10^. For 454 sequencing, we adjust the indel quality score of the *j*-th nucleotide position in a homopolymer run starting from the second nucleotide in the repeat, *q*_*i*_ =*max*(*q*-*j*\*5, 10), i.e., the quality scores degrade linearly, till a threshold maximum error, beyond which it is maintained constant, and leave *q*_*s*_ intact. This dual quality score also comes handy for handling “N”-nucleotides. The “N”-nucleotide normally indicates the existence of a nucleotide, but with a very low confidence to discern which nucleotide it actually is. For such positions, we adjust *q*_*i*_ to be 20 (low probability 1% of making an indel) and leave *q*_*s*_ unchanged (at a high probability corresponding to the low quality score).

### Statistical Significance of Meta-Align

The probability value output from Meta-Align is an indication of the sequence similarity between the input pair of reads. To evaluate the statistical significance of the probability value found by Meta-Align, we follow the Z-value approach first proposed by [24] and extensively researched by [25]. We consider a pair of sequences (*X, Y*) and obtain an alignment score *S(X, Y)* using Meta-Align. We randomly shuffle one of the sequences, *Y* to *Y\**, and compute the score *S(X, Y*)* using Meta-Align. By performing this shuffling 250 times we can obtain a Z-statistics,

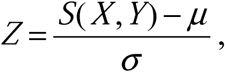

where μ and σ are the mean and the standard deviation of scores *S(X, Y*)* from random shuffling.

As shown in Figure 5, a Z-score of 11.23 was obtained between a pair of sequences from BAliBASE based on 250 random shuffles. It has been proposed by [25], based on Chebyshev inequality, that a Z score about 8 is an indicator that the alignment scores are significant. Therefore, we conclude that the alignment generated by Meta-Align is statistically significant. Actually, it can be seen from Figure 5 that the score distribution of Meta-Align has an exponential-like tail which may relate to the Gumbel distribution of Viterbi scores [26-28], and this Z-score of 11.23 is probably more significant than what Chebyshev inequality suggested.

**Figure 5.**
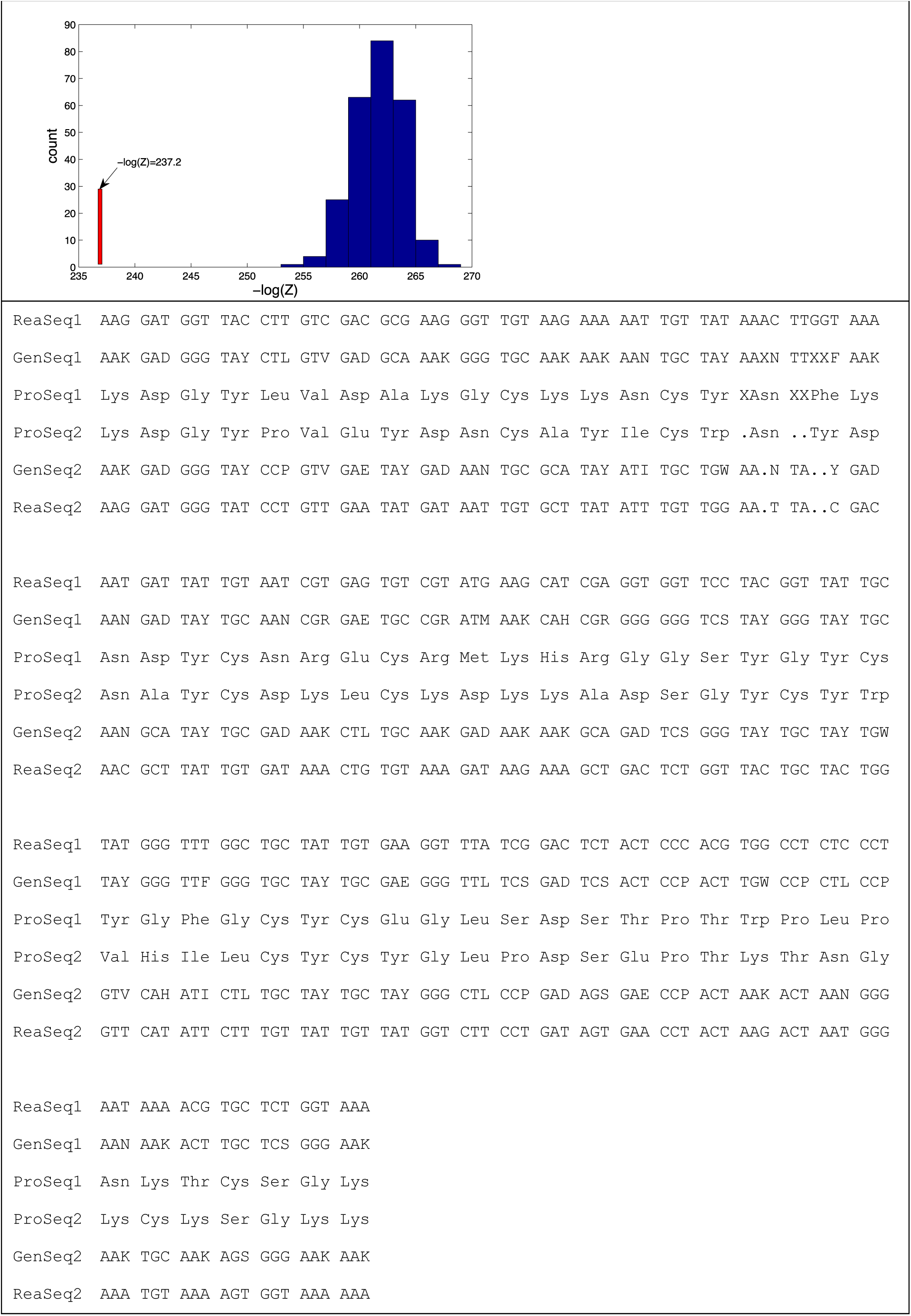
Statistical significance of the Meta-Align’s score of two sequences from BAliBASE. Panel (a) shows the distribution of Meta-Align scores for 250 randomly shuffled sequences in blue, and the score for the pair of real sequence in red. Panel (b) shows Meta-Align’s output for the pair of sequences in 6-tract format: Each block representing a CMU match, tract 1 and track 6 represent the input pair of reads. Tract 2 and tract 5 shows the inferred error-corrected DNA sequences of the input pair, where codon AAK represents a degenerated CMU matching AAX where the symbol K in the last position represents the amino acid encoded by the degenerated codon. Tract 3 and tract 4 represent the amino acid match.

## Discussion

Metagenomic sequence data differs from traditional sequence data in a number of important ways. Most importantly, most microbial communities are diverse and highly uneven. Consequently, and the amount of sequence data that is tractable to acquire is a tiny fraction of the total genetic material in the community. As a result, sequence reads in metagenomic sequencing projects sample most underlying genomes only sparsely, and so many reads cannot be assembled (except perhaps reads from a few dominant species). Protein-coding regions found on these reads are consequently highly fragmentary; due to the random nature and limited read length of sequencing, a sequenced read typically overlaps only part of the complete coding sequence in the original genome from which it is derived. In the case of isolate genome sequencing, high coverage allows detection and correction of sequencing errors; in contrast, these errors typically remain uncorrected in the metagenomic case, where (even high quality) reads are mostly single-pass and so cannot be confirmed. To date, scant attention has been given to the impact of fragmentation and errors on downstream data analysis.

We addressed the issues in the present study. Our approach has a key innovation to incorporate quality scores from sequencing and also to accommodate the uncertainty created by gaps in or between codons. As a result, our approach is capable of accounting for all types of errors of NGS data. Specifically, in our algorithm each triplet (possible codon) is not assigned a single amino acid translation as would ordinarily be the case. Instead, each possible codon is treated as a distribution of amino acids, in which each of the 20 possible amino acids is assigned a probability of being encoded by the combination of DNA bases and gaps contained within the codon. To construct distributions for each possible codon from a DNA sequence, we begin with the quality score for each called DNA base, which determines the probability of the called base; the remainder of the probability is distributed equally among the other three potential nucleotides.

Indel errors are frequently associated with homopolymers, particularly in the case of pyrosequencing. While we could include details of this tendency in the algorithm, we currently believe it will be sufficient to apply a uniform error rate. We will compute realistic aggregate indel rates based on the distribution of species with different GC contents observed in real metagenomic data sets. This will enhance the accuracy of Meta-Align.

While, at present, this approach is not fast enough for large-scale database searches, we think that this will not be a concern because majority of data can be processed commonly available tools. Making more realistic models by including more precise information will enhance our algorithm. We will also develop a version of the algorithm, such as BLASTX, to permit nucleotide versus protein alignments. This will often be the case when metagenomic data are aligned with sequences from protein databases. We believe our algorithm can be also highly useful for this purpose, in addition to pairwise gene finding.

## Conclusions

We have developed a novel HMM-based algorithm, Meta-Align that allows for aligning DNA sequences in protein space and incorporating quality scores to address the issues caused by insertions and deletions which are relatively common in metagenomic data. Meta-Align is capable of accounting for the frameshifts, substitutions, and indels, and obtaining the most likely alignment of the amino acids produced by matching pairs of triplets, that are predicted codons in each source genomic sequence. Each triplet may contain any numbers of gaps in its DNA sequence, thereby seamlessly accommodating frameshift errors into the alignment process. We have shown that Meta-Align outperforms TBLASTX using a diverse set of simulated reads, and that Meta-Align can produce better alignments using real quality scores. Thus, we expect that Meta-Align will be particularly useful when applied to error-prone read sequences with low quality scores.

## Availability and requirements

Project name: Metaalign

Project home page: https://github.com/shravan-repos/Metaalign

Operating system(s): Linux

Programming language: C++

License: GNU GPL

## Authors’ contributions

KT and DZ jointly designed the algorithm. KT, SK, and DZ implemented the algorithm, and drafted the manuscript. SK and DZ conducted the benchmarking tests. SEB contributed to the conception of the project and edited the manuscript. All authors approved the final manuscript, though SEB was unable to review the final manuscript due to his injury.

## Acknowledgements

The authors would like to thank David Soergel, Robert C. Edgar, and Daniel Richter for valuable discussions. This work is partly funded by NIH grant R00 RR024163. SK was partly funded by AU EPSCoR NSF-funded Graduate Research Scholars Program.

